# Transcriptional regulation of MGE progenitor proliferation by PRDM16 controls cortical GABAergic interneuron production

**DOI:** 10.1101/2019.12.17.879510

**Authors:** Miguel Turrero García, José-Manuel Baizabal, Diana Tran, Rui Peixoto, Wengang Wang, Yajun Xie, Manal A. Adam, Salvador Brito, Matthew A. Booker, Michael Y. Tolstorukov, Corey C. Harwell

**Author notes:** Department of Biology, Indiana University, Bloomington IN 47405, USA. Department of Psychiatry, University of Pittsburgh, Pittsburgh PA 15219, USA.

## Abstract

The mammalian cortex is populated by neurons derived from neural progenitors located throughout the embryonic telencephalon. Excitatory neurons are derived from progenitors located in the dorsal telencephalon, while inhibitory interneurons are generated by ventral telencephalic progenitors. The transcriptional regulator PRDM16 is expressed by radial glia, neural progenitors present in both regions; however, its mechanisms of action are still not fully understood. It is unclear if PRDM16 functions plays a role in neurogenesis in both dorsal and ventral progenitor lineages, and if so, whether it does so by regulating common or unique networks of genes. Here, we show that *Prdm16* expression in MGE progenitors is required for maintaining their proliferative capacity and for the production of proper numbers of pallial GABAergic interneurons. PRDM16 binds to cis-regulatory elements and represses the expression of region-specific neuronal differentiation genes, thereby controlling the timing of neuronal maturation. Our results highlight the importance of PRDM16 for the development of both excitatory and inhibitory cortical circuits. We demonstrate the existence of convergent developmental gene expression programs regulated by PRDM16, which utilize both common and region-specific sets of genes in the cortex and the MGE to control the proliferative capacity of neural progenitors, ensuring the generation of correct numbers of cortical neurons.

## INTRODUCTION

The complex circuitry of the mammalian neocortex comprises two major types of neuronal cells, excitatory pyramidal neurons and inhibitory interneurons, with different functions and origins (Campbell, 2003; Wilson and Rubenstein, 2000). In the mouse, both types of neurons are generated between embryonic day (E)10 and E16, from neural progenitors residing in the proliferative zones located along the lateral ventricles of the telencephalon (Anthony et al., 2004; Marin and Muller, 2014). Excitatory neurons are produced from dorsal (pallial) telencephalic proliferative zones (Govindan and Jabaudon, 2017), while the majority of inhibitory interneurons are produced from two transient proliferative structures in the ventral (subpallial) telencephalon, known as the medial (MGE) and caudal (CGE) ganglionic eminences (Miyoshi et al., 2010; Nery et al., 2002; Wonders and Anderson, 2006; Xu et al., 2004). Progenitors in the MGE give rise to two major groups of inhibitory interneurons, fast-spiking and non-fast-spiking, which can be identified by their expression of the markers parvalbumin (PV) and somatostatin (SST), respectively (Xu et al., 2004). The majority of SST-expressing interneurons are born in the early stages of neurogenesis, while PV-expressing cells are generated later (Hu et al., 2017; Miyoshi et al., 2007). Both populations undergo a process of tangential migration into the developing cortex, following a spatiotemporal pattern in which earlier born cells typically occupy the deep layers of the cortex, while later born cells occupy more superficial layers after their tangential migration (Lopez-Bendito et al., 2004).

During neurogenesis, neural progenitors in the MGE transition through a series of competence states as they produce these two types of interneurons: radial glia (RG) divide at the surface of the ventricle, self-renewing while giving rise to either neurons (direct neurogenesis) or more committed transit-amplifying progenitors with limited self-renewal capacity, that usually divide once to generate two neurons (indirect neurogenesis) (Harwell et al., 2015; Turrero Garcia and Harwell, 2017). In order to understand how the balance of excitation and inhibition is achieved in the developing cortex, it is necessary to know the mechanisms by which MGE progenitors regulate their proliferation and consequently their neuronal output (Lim et al., 2018; Petryniak et al., 2007). One critical factor that ensures correct neural progenitor amplification and cell lineage progression is PRDM16, a transcriptional regulator that is specifically expressed in radial glia of the developing telencephalon (Baizabal et al., 2018; Chuikov et al., 2010; Shimada et al., 2017). In the cortex, *Prdm16* controls indirect neurogenic divisions, radial glia lineage progression and the production of late born upper layer pyramidal neurons by regulating the epigenetic state of developmental enhancers (Baizabal et al., 2018). PRDM16 is also expressed in radial glia of the MGE, but it is unknown whether it regulates lineage progression programs controlling the production of cortical interneurons. Here we show that *Prdm16* expression in MGE progenitors is necessary to maintain their proliferative potential and ensure the generation of sufficient numbers of cortical interneurons and the maintenance of correct inhibitory input onto pyramidal cells. PRDM16 controls the expression of both generic and MGE-specific neuronal differentiation programs through its binding to sets of cis-regulatory elements that are either common between this area and the developing neocortex or exclusive to the MGE.

## RESULTS

### *Prdm16* expression in MGE progenitors regulates the number of cortical interneurons

We generated a Cre-lox system-based conditional knockout mouse model, in which *Prdm16* was deleted in cells with a developmental history of *Nkx2.1* expression (i.e., neural progenitors in the MGE and their progeny) (Cohen et al., 2014; Xu et al., 2008). Additionally, these mice carried a Cre-dependent fluorescent reporter, tdTomato (tdT), that allows identification of cortical interneurons derived from *Nkx2.1*-expressing MGE progenitors in the adult brain, after the expression of this gene is shut down (**Figures 1A, 1C**) (Madisen et al., 2010). In order to verify the specific ablation of *Prdm16* in *Nkx2.1*-expressing cells, we examined the brains of wildtype (WT) and knockout (KO) mice at embryonic day (E)13, a developmental stage midway through cortical interneuron neurogenesis (Turrero Garcia and Harwell, 2017), confirming the absence of PRDM16 in the MGE by immunostaining (**Figure 1B**). We analyzed the brains of postnatal day (P)30 mice, in order to understand the consequences of *Prdm16* deletion in the mature cortex. We performed immunofluorescence staining to detect tdT, somatostatin (SST), and parvalbumin (PV), markers of the two major subgroups of MGE-derived forebrain interneurons (**Figure 1C**). We found that the number of tdT-positive (tdT+) interneurons in KO cortices was 27.27 % lower than in WT controls (**Figure 1D**). Decreased numbers of tdT+ cells were observed across all cortical layers except for layer VI (**Figure 1E**), and the proportion of tdT+ cells that were positive for either SST or PV was the same in WT (27.39 % SST+ and 29.96 % PV+) and KO cortices (26.60 % SST+ and 27.30 % PV+). The total numbers of tdT+ cortical interneurons expressing SST (**Figure 1F**) or PV (**Figure 1H**) were decreased by similar proportions as the overall tdT+ cells (30.43 % in SST+ and 35.03 % in PV+ cells). The loss of SST+ (**Figure 1G**) and PV+ (**Figure 1I**) tdT+ cortical interneurons was also observed across all cortical layers except for layer VI. We observed a similar phenotype in the hippocampus (**Figure S1A**), where there was a decrease in the number of tdT+ *Nkx2.1*-lineage interneurons (**Figure S1B**), which was consistent across all regions except the subiculum (**Figure S1C**), and across all layers (**Figure S1D**). In the hippocampus, the loss was cell type-specific: the number of SST+, tdT+ cells was decreased (**Figure S1E**); unexpectedly, PV+, tdT+ cells were not affected (**Figure S1F**). As in the cortex, the distribution of tdT+ cells in the hippocampus of KO mice closely recapitulated that of the WT in all areas (**Figures S1G, S1H**). Together, these observations indicate that *Prdm16* regulates the numbers of MGE-derived interneurons in the adult cortex and hippocampus.

**Figure 1:**
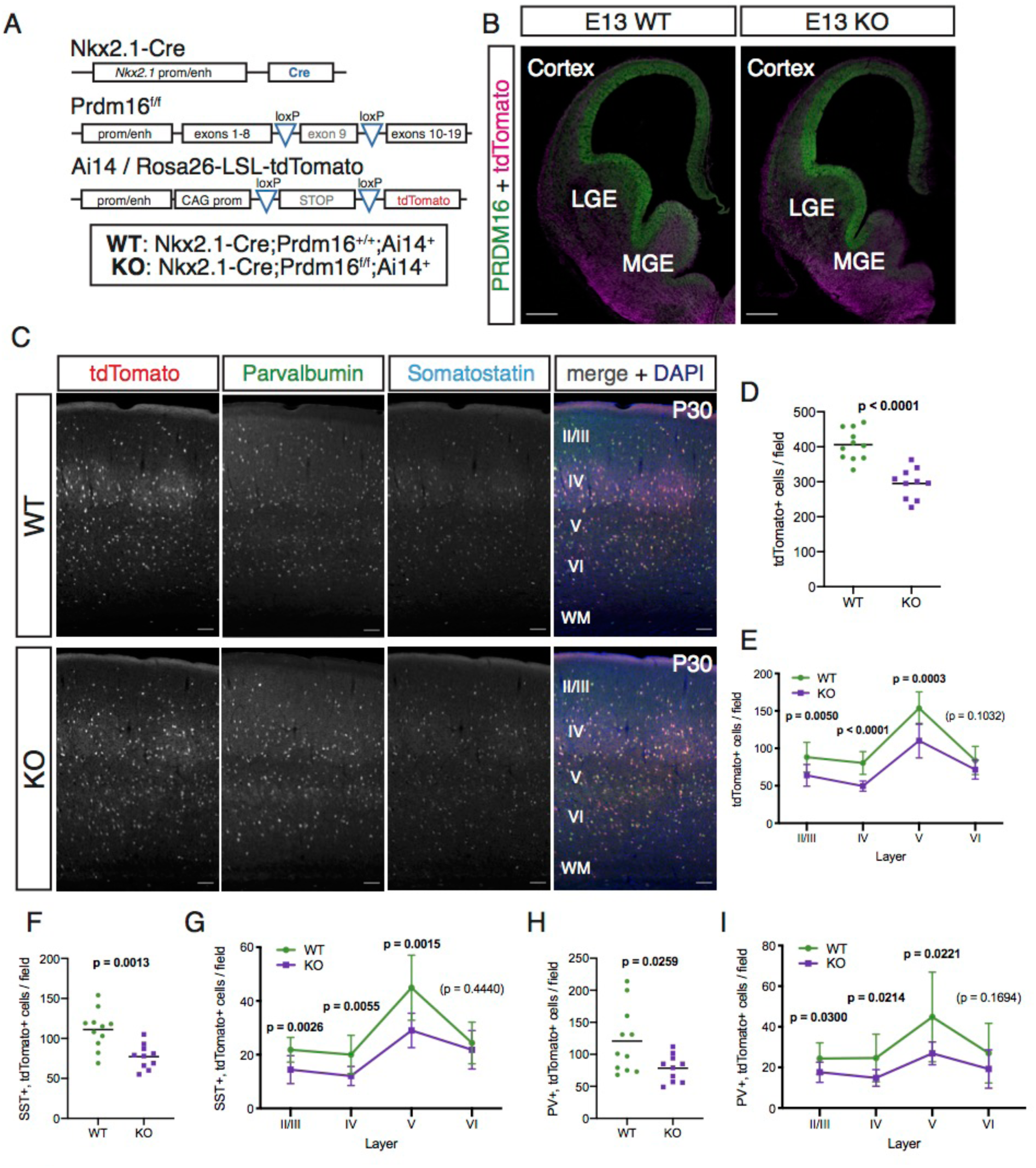
Deletion of *Prdm16* in the *Nkx2.1* lineage causes a loss of cortical interneurons. **A)** Schematic of genetic strategy for conditional deletion of Prdm16 and simultaneous fate mapping of Nkx2.1 expressing cells. **B)** Immunostaining for PRDM16 (green) and tdTomato (magenta) on forebrain coronal sections of embryonic day (E)13 WT and KO mice, showing the loss of PRDM16 in the ventricular zone of the *Nkx2.1*-expressing medial ganglionic eminence (MGE), but not in the lateral ganglionic eminence (LGE) or cortex. Scale bars: 250 *µ*m. **C)** Immunostaining for tdTomato (red in merged image), parvalbumin (green in merge) and somatostatin (cyan in merge), counterstained with DAPI (blue in merge), in the cortex of postnatal day (P)30 WT and KO mice. Cortical layers are indicated in white. Scale bar, 100 *µ*m. **D)** Total number of tdTomato+ cells per 1 mm-wide column spanning the entirety of the cortex. **E)** Number of tdTomato+ cells in each indicated cortical layer, quantified per 1 mm-wide column. **F)** Total number of cells co-labeled with somatostatin and tdTomato (SST+, tdTomato+) per 1 mm-wide cortical column. **G)** Number of SST+, tdTomato+ cells in each indicated cortical layer, per 1 mm-wide column. **H)** Total number of cells co-labeled with parvalbumin and tdTomato (PV+, tdTomato+) per 1 mm-wide cortical column. **I)** Number of PV+, tdTomato+ cells in each indicated cortical layer, per 1 mm-wide column. Analysis in panels **C-J** was performed on samples from P30 WT (green circles) and KO (purple squares) animals. Black bars in panels **D**, **F** and **H** indicate the mean. Mean ± S.D. is displayed in panels **E**, **G** and **I**. Unpaired t-tests with Welch’s correction (panels **D**, **F**, **H**) or multiple t-tests (panels **E**, **G**, **I**) were performed; p-values are indicated above the corresponding compared sets of data: those highlighted in bold represent significant differences (p<0.05).

### Partial compensation for loss of MGE-derived cortical interneurons in the KO cortex

The proper complement of inhibitory interneurons is essential for maintaining the excitatory and inhibitory network balance; however, KO mice did not display obvious seizure activity (Neves et al., 2013; Southwell et al., 2014). We hypothesized that loss of MGE-derived interneurons could be compensated by populations derived from the other major embryonic source of cortical inhibitory neurons, the caudal ganglionic eminence (CGE) (Denaxa et al., 2018). To test this, we performed immunofluorescence staining in P30 WT and KO brains to detect reelin and vasointestinal peptide (VIP), two markers of CGE-derived cortical interneurons (**Figure 2A**) (Lee et al., 2010; Miyoshi et al., 2010). We did not detect a significant difference in the total number of VIP+ (**Figure 2B**) or reelin+ (**Figure 2C**) cortical interneurons. Their distribution across cortical layers I-VI was largely similar between WT and KO animals (**Figures S2A, S2B**); however, there was a significant increase in the number of reelin+ cells in layer I of the mutant cortex (**Figure S2B**). Reelin+ interneurons originate from either the CGE or the MGE (Miyoshi et al., 2010); the latter population is largely SST+ (Pesold et al., 1999). To distinguish between these two reelin+ subpopulations, we quantified the proportion of cortical reelin+ cells that were tdTomato+, since cells labeled with the Nkx2.1-Cre driver line are originated in the MGE. We found that approximately two thirds (67.16 %) of reelin+ interneurons in WT cortices were MGE-derived, but this proportion was significantly lower (52.41%) in KO mice (**Figure 2D**). KO cortices contained an increased number of CGE-derived (i.e. tdTomato–) reelin+ interneurons (**Figure 2E**), concentrated in cortical layers I and II-III (**Figure 2F**). We did not observe an analogous increase in the VIP+ population (**Figure S2A**). These results suggest that there is an increase in the number of CGE-derived interneurons in the upper layers of KO cortices, which could partially compensate for the decrease in the number of MGE-derived cortical interneurons. This effect was specific to reelin+ interneurons, which express a common marker while being derived from different developmental origins. No increase was observed in VIP+ interneurons, which is consistent with evidence that survival and maturation of this subgroup is not dependent upon activity (**Figure S2C**) (De Marco García et al., 2011; Priya et al., 2018).

**Figure 2:**
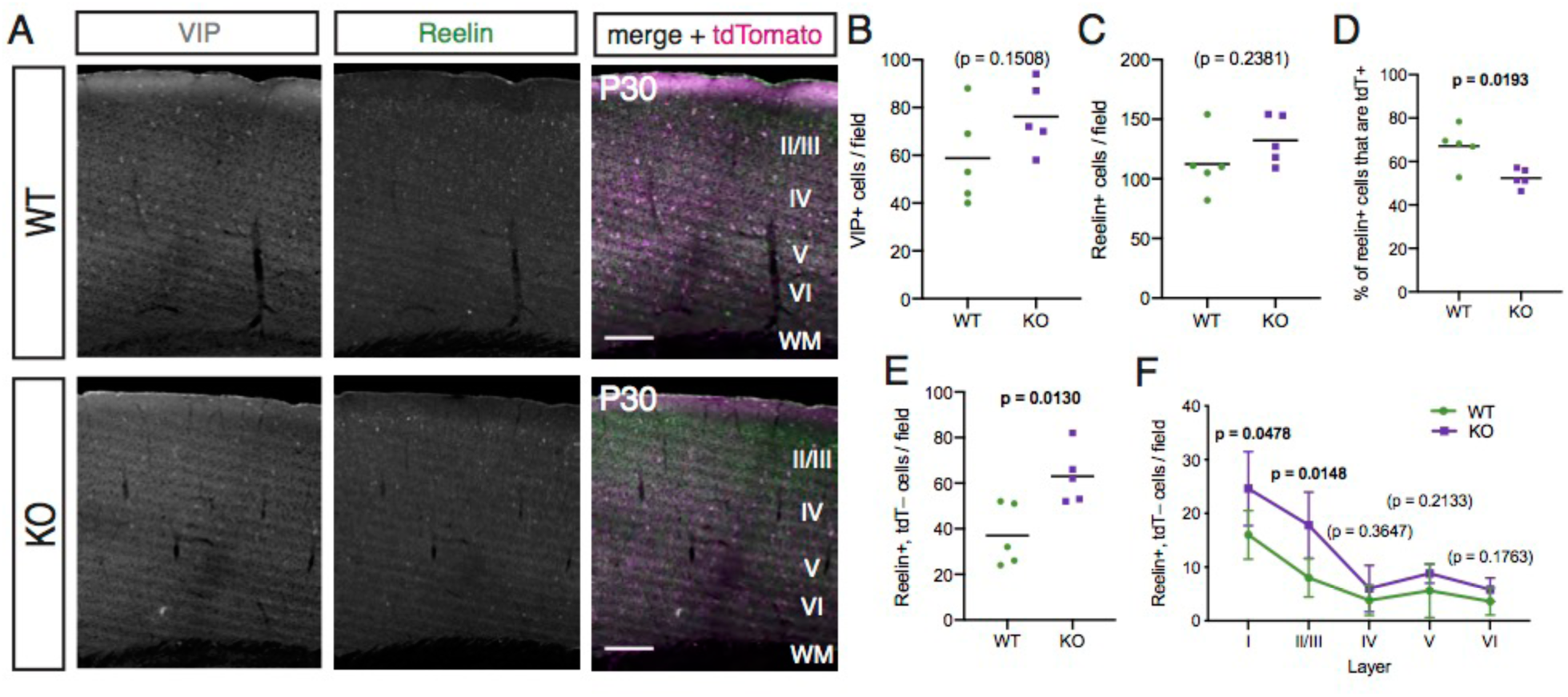
The loss of *Nkx2.1*-lineage cortical interneurons in *Prdm16* mutants is partially compensated by an increase in non-MGE-derived reelin+ cells in upper layers. **A)** Immunostaining for vasointestinal peptide (VIP, green in merge), reelin (gray in merge), and tdTomato (magenta in merge) in the cortex of P30 WT and KO mice. Cortical layers are indicated in white. Scale bar, 200 *µ*m. **B)** Total number of VIP+ cells per 1 mm-wide column spanning the entirety of the cortex. **C)** Total number of reelin+ cells per 1 mm-wide cortical column. **D)** Percentage of reelin+ cells costained for tdTomato. **E)** Total number of reelin+, tdTomato– cells per 1 mm-wide column. **F)** Number of reelin+, tdTomato– cells in each indicated cortical layer, per 1 mm-wide column. Analysis in panels **A-F** was performed on samples from P30 WT (green circles) and KO (purple squares) animals. Black bars in panels **B**-**E** indicate the mean. Mean ± S.D. is displayed in panel **F**. Unpaired t-tests with Welch’s correction (panels **B**-**E**) or multiple t-tests (panel **F**) were performed; p-values are indicated above the corresponding compared sets of data: those highlighted in bold represent significant differences (p<0.05).

### Decreased inhibitory input onto pyramidal neurons in the KO cortex

We performed whole cell electrophysiological recordings to investigate whether the reduction in number of interneurons that we observed in the cortex of adult KO mice led to a decrease of inhibitory input onto pyramidal cells. We recorded from excitatory neurons in layers II/III of the cortex of three weeks old WT and KO mice, analyzing their inhibitory input by comparing the frequency and amplitude of spontaneous miniature inhibitory postsynaptic currents (mIPSC; **Figure 3A**). We found that the frequency of mIPSCs was 33.54% lower in the KO cortex (**Figure 3B**), while their amplitude was not significantly different (**Figure 3C**), consistent with an overall decrease in the number of inhibitory GABAergic inputs. From this, we conclude that the loss of MGE-derived interneurons in the cortex of KO mice results in a decrease in the inhibitory input to pyramidal neurons in the upper layers that cannot be fully rescued by the increase in the number of upper layer CGE-derived reelin+ interneurons.

**Figure 3:**
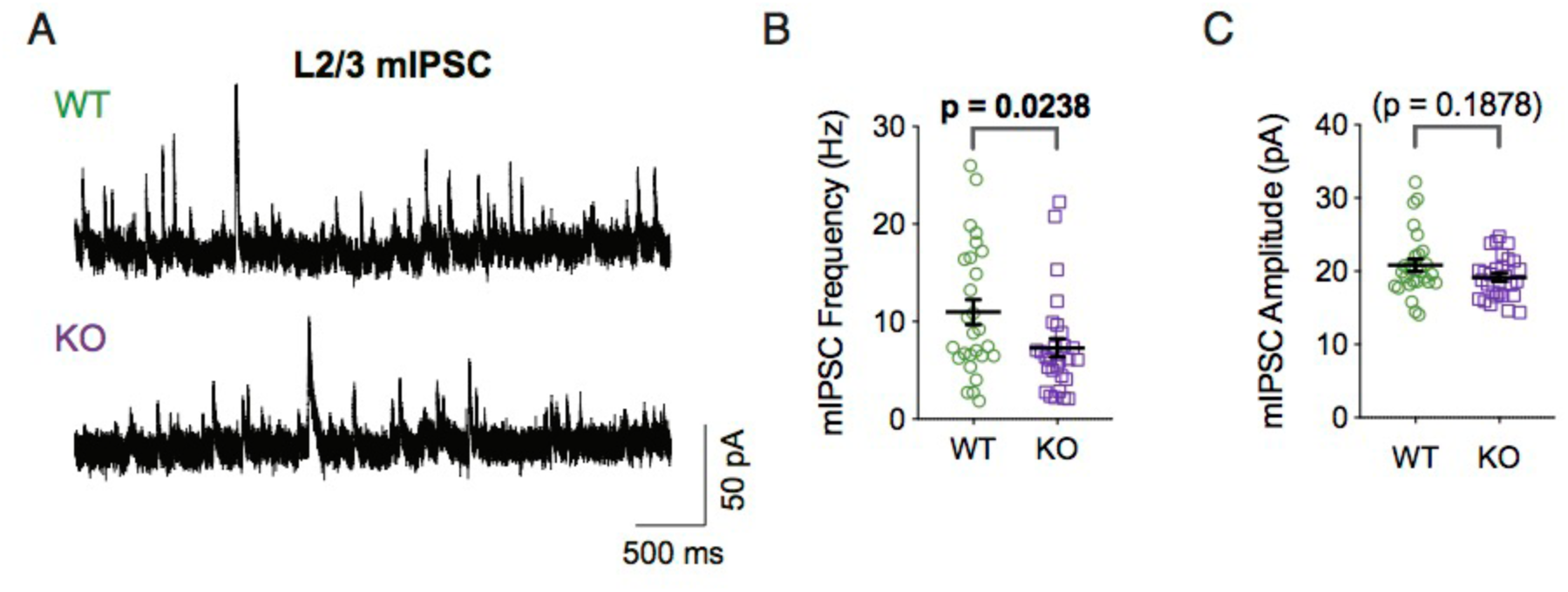
Pyramidal neurons in the cortex of *Prdm16* mutant mice receive decreased inhibitory inputs. **A)** Representative traces of miniature inhibitory postsynaptic currents (mIPSCs) recorded from layer II/III pyramidal cells of the somatosensory cortex of P21 WT and KO mice. **B**, **C)** Frequency (in Hertz, **B**) and amplitude (in pA, **C**) values of mIPSCs. N=3 mice, n=27 cells (WT), N=3 mice, n=28 cells (KO). All values are displayed, with mean ± S.E.M. indicated in black. Mann-Whitney tests (panels **I**, **J**) were performed; p-values are indicated above the corresponding compared sets of data: the one highlighted in bold represents a significant difference (p<0.05).

### *Prdm16* regulates MGE progenitor proliferation

We sought to understand the causes leading to the decreased numbers of tdT+ cortical interneurons in the cortex of KO mice. One possibility is that loss of *Prdm16* function in MGE-derived postmitotic neurons could decrease their survival (Close et al., 2012; Denaxa et al., 2012). However, immunostaining for the apoptosis marker cleaved caspase-3 did not differ between WT and KO cortices at any of the developmental stages analyzed (**Figure S3A**). Another possibility is that defects in the migratory capacity of *Nkx2.1*-lineage interneurons could impair their ability to reach their final destination in the cortex through tangential migration (Anderson et al., 1997; Corbin et al., 2001). We examined the position of tdTomato+ interneurons in the developing brains of WT and KO animals, and found no differences between them (**Figure S3B**). We concluded that *Prdm16* must exert its role at the progenitor level, guiding the generation of cortical interneurons rather than their survival or long-distance migration.

We analyzed the *Nkx2.1*-expressing progenitors that give rise to these cells. First, we compared the number of progenitors in the MGE of WT and KO mice at E13, a mid-neurogenic stage (Turrero Garcia and Harwell, 2017). We performed immunofluorescence staining for phosphorylated histone 3 (pH3), a marker of late G2/M-phase in cycling cells (**Figure 4A**). We found a significant reduction in the overall density of dividing cells in the MGE of KO mice (**Figure 4B**); this effect could be observed both in the ventricular zone (VZ), where radial glia reside (**Figure 4C**), and in the subventricular zone (SVZ) (**Figure 4D**), which harbors other types of transit-amplifying progenitors (Turrero Garcia and Harwell, 2017). In order to assess the proliferative capacity of MGE progenitors, we infected embryonic mice with GFP-expressing retrovirus, which allowed us to identify and analyze the progeny (or “clones”) derived from single radial glial cells 24 h after infection (**Figure 4E**) (Baizabal et al., 2018; Harwell et al., 2015). While the majority of clones in the MGE of control mice were composed of two cells, with a smaller proportion of 1-cell clones and progressively smaller proportions of bigger clones, in the MGE of KO mice this distribution was shifted towards smaller clone sizes: almost two thirds of the clones were composed of a single cell, and about one third of the total were two-cell clones (**Figure 4F**). Correspondingly, the average clone size was smaller in the MGE of KO mice (**Figure 4G**). These results indicate that the changes in cortical interneuron output that we observed are due to defects in the proliferative capacity of MGE progenitors.

**Figure 4:**
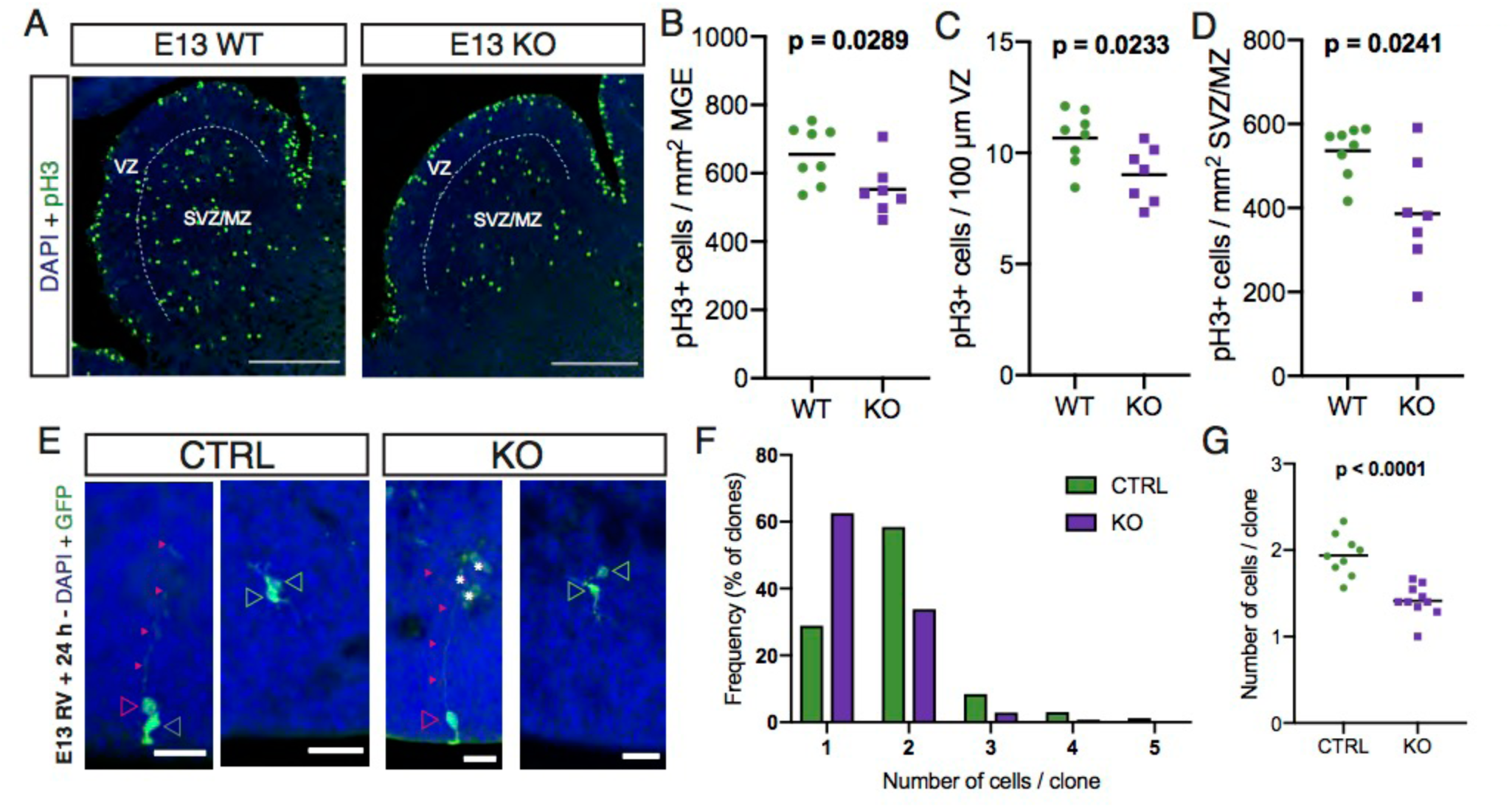
The loss of cortical interneurons in the KO cortex is caused by defects in MGE progenitor proliferation. **A)** Representative images of immunofluorescence experiments performed on the MGE of WT and KO mice at E13, stained for the mitotic marker pH3 (green), and tdTomato (magenta); nuclei were counterstained with DAPI (blue). Scale bars, 200 *µ*m. The ventricular (VZ) and subventricular/mantle (SVZ/MZ) zones are indicated. **B)** Quantification of the number of pH3+ cells per mm^2^ in the MGE of WT (green circles) and KO (purple squares) mice at E13. **C)** Quantification of the number of VZ pH3+ cells per 100 *µ*m of ventricular surface length in the MGE of WT (green circles) and KO (purple squares) mice at E13. **D)** Quantification of the number of pH3+ cells per mm^2^ in the SVZ/MZ of the MGE of WT (green circles) and KO (purple squares) mice at E13. **E)** Representative images of retrovirus-labeled clones in the MGE of WT (left) and KO (right) mice, 24 h after infection. Clones were typically composed of radial glia (magenta empty arrowheads), distinguished by their long basal processes (small magenta arrowheads) sometimes contacting blood vessels (white asterisks), and/or intermediate progenitors (green empty arrowheads) or other unidentified cells (gray empty arrowhead). Scale bars, 25 *µ*m. **E)** Frequency distribution of retrovirally-labeled clone size in the MGE of WT (green; n = 165 clones from 9 embryos) and KO (purple; n = 141 clones from 10 embryos) mice at E14, 24 h after infection. **F)** Number of cells per MGE clone of WT (green circles) and KO (purple squares) E14 mice, 24 h after retroviral infection. The average clone sizes for each sample (n = 9 for WT, n = 10 for KO) are represented. Black bars in panels **C** and **F** indicate the mean. Unpaired t-tests with Welch’s correction (panels **B**-**D**, **G**) were performed; p-values are indicated above the corresponding compared sets of data: those highlighted in bold represent significant differences (p<0.05).

### PRDM16 associates with cis-regulatory elements involved in nervous system development

In the cortex, *Prdm16* regulates neurogenesis by modulating the expression of genes involved in the generation of intermediate progenitors and neuronal differentiation (Baizabal et al., 2018). To understand the molecular mechanisms by which *Prdm16* regulates the proliferation and neuronal output of MGE progenitors, we performed ChIP-Seq for PRDM16 on MGE samples isolated from E14 wildtype mice. We detected 3517 statistically reproducible (IDR p<0.05) ChIP-Seq peaks across two experimental replicates, representing PRDM16 binding sites. We compared the enrichment of PRDM16 binding sites with those previously characterized in PRDM16 ChIP-Seq experiments performed in E15 cortex (Baizabal et al. 2017) and histone modifications in the MGE at E13 obtained from a published dataset (Sandberg et al. 2016). We found that potential PRDM16 binding sites largely overlap between the MGE and cortex datasets, and they are enriched in open chromatin histone marks H3K4me3, H3K4me1, H3K27ac, but not in the repressive H3K27me3 modification (**Figures 5A, B**). The distribution of PRDM16 ChIP-Seq peaks was similar to that of the E15 cortex dataset (**Figure S5A**) (Baizabal et al. 2017). We analyzed the distribution of PRDM16 peaks across the genome, and found that the majority of peaks are located in intergenic and intronic genomic regions. About 10 % of PRDM16 binding peaks were associated with transcription start sites (TSS) (**Figure 5C**). We then examined the genes closest to MGE PRDM16 peaks by performing gene ontology (GO) analysis. The nearest genes in every annotated genomic region were associated to similar GO terms, with a relative enrichment of terms related to DNA synthesis and repair in genes close to TSS-associated peaks. (**Figure S4B**). All of the top 30 GO terms for genes were related to developmental processes, including the generation and migration of neurons (**Figure 5D**). We compared the MGE and cortex ChIP-Seq datasets in order to understand which PRDM16 peaks were exclusive to MGE progenitors. We generated a list of highly enriched peaks (>3-fold enrichment over input) for both datasets and obtained a list of 3206 peaks. About one third of these were MGE-exclusive, while two thirds were common to both datasets, and less than 10 % were cortex-specific (**Figure 5E**). GO terms of nearest genes for both subsets of peaks had overall similar representation, with a slightly lower representation of synaptic signaling-related terms in MGE-exclusive peaks (**Figure S5B**). It is possible that PRDM16 may exhibit lineage-specific genomic binding in order to direct GABAergic versus glutamatergic cell fate specification. To explore this possibility, we analyzed the sequence of PRDM16 peaks searching for *de novo* or known binding motifs either in all MGE peaks (**Figure 5F**) or in the MGE-specific subset (**Figure 5G**). MGE peaks that overlapped with cortical peaks were enriched in binding motifs for transcriptional regulators associated with neurogenesis, including LHX2 and SOX10 (Chou and Tole, 2019; Weider and Wegner, 2017). MGE-exclusive peaks showed motif enrichment for TCF12 and ASCL1, bHLH transcription factors involved in progenitor proliferation and neuronal cell fate determination (Dennis et al., 2019). All together these data suggest that PRDM16 may form MGE-specific transcription factor complexes to regulate transcriptional programs guiding the specification of cortical interneurons (Sandberg et al., 2018).

**Figure 5:**
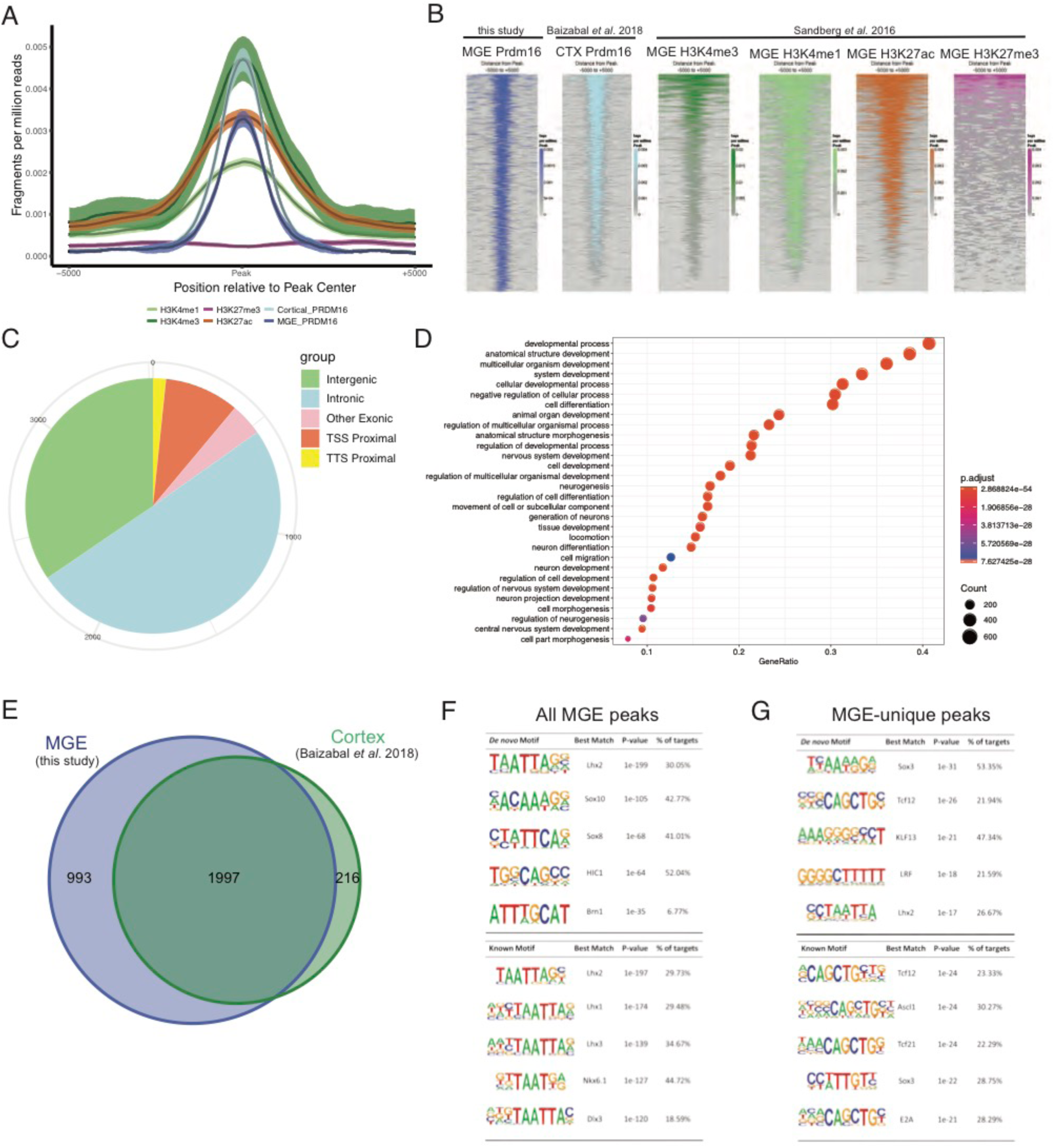
PRDM16 binding sites in the MGE are associated with open chromatin marks and genes implicated in neural development. **A)** Read density (fragments per million reads) in several embryonic ChIP-Seq experiments (color-coded as in and explained in **B**), within a genomic window centered around PRDM16 binding sites in the embryonic cortex (light blue) or the MGE (all other plots). **B)** read density alignment of the ChIP-Seq experiments depicted in **A**; data were generated for this study from E14 MGE (dark blue), or obtained from Baizabal *et al*. 2017 (CTX, ChIP-Seq for PRDM16 in E15 cortex; light blue) and Sandberg *et al*. 2016 (ChIP-Seq for each specified histone modification in E13 MGE; all other plots). **C)** Distribution of E13 MGE PRDM16 ChIP-Seq peaks across the genome (TSS: transcription start site; TTS: transcription termination site). **D)** Gene ontology term enrichment in potential PRDM16 binding sites in the MGE, obtained by analysis of genes closest to ChIP-Seq peaks. **E)** Overlap of PRDM16 peaks between cortex and MGE ChIP-Seq experiments. The number of peaks within each sector of the Venn diagram is indicated. **F, G)** *De novo* (top) and known (bottom) motif analysis of PRDM16 ChIP-Seq peaks, in all MGE peaks (**F**) or MGE-exclusive peaks (**G**).

### PRDM16 downregulates neuronal differentiation genes

In order to understand the function of PRDM16 *in vivo*, we selected two genes known to play a role in neuronal differentiation and function: *Pdzrn3* and *Gad2*. Several ChIP-Seq peaks in the *Pdzrn3* genomic locus were common to the MGE and cortex datasets (**Figure 6A**). We previously showed that *Pdzrn3* repression by PRDM16 controls the migration of pyramidal neurons in the cortex (Baizabal et al. 2017). A ChIP-Seq peak in the *Gad2* genomic locus was exclusive to the MGE (**Figure 6B**). GAD2 is required for GABA synthesis, an essential feature of inhibitory interneuron function. In order to understand how PRDM16 binding controls cortical interneuron specification through *Pdzrn3* and *Gad2*, we compared the expression levels of these genes in the MGE of WT and KO mice at E13. We first performed RT-PCR experiments on MGE tissue from embryos of either genotype, and found a consistent increase in the expression levels of both *Pdzrn3* and *Gad2* in KO tissue, suggesting that PRDM16 directly represses the expression of these genes to prevent premature differentiation of MGE progenitors (**Figure 6C**). In order to directly visualize the spatial pattern of *Pdzrn3* and *Gad2* expression we used single molecule fluorescent *in situ* hybridization (RNAscope) on embryonic tissue sections of WT and KO embryos. We detected RNA puncta representing single mRNA transcripts of either *Pdzrn3* (**Figure 6D**) or *Gad2* (**Figure 6F**). Automated quantification of these puncta in the VZ and SVZ/MZ revealed that there was a significant increase in the expression levels of both genes in KO animals that was specific to the VZ in both cases (**Figures 6E, G**). This suggests that PRDM16 primarily acts in VZ radial glia to repress the expression of genes involved in neuronal differentiation, thus allowing these cells to maintain their proliferative capacity and transition through transit-amplifying stages in order to generate sufficient numbers of cortical interneurons.

**Figure 6:**
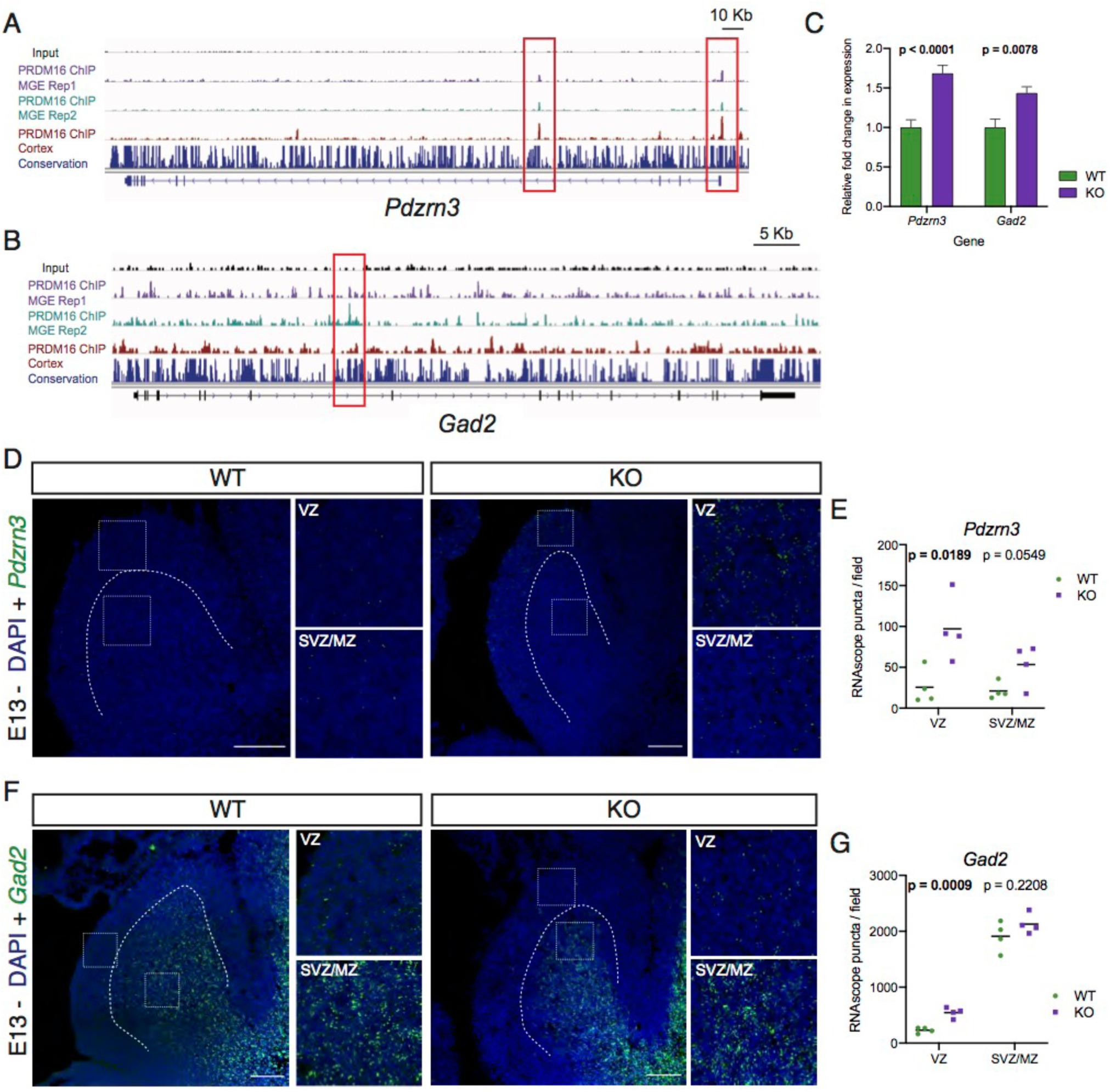
PRDM16 represses the expression of genes involved in neuronal differentiation in the MGE. **A, B)** ChIP-Seq tracks showing PRDM16 binding sites in two ChIP-Seq experimental replicates in the E14 MGE (middle tracks), compared to E15 cortex (bottom track), in the loci of *Pdzrn3* (**A**) and *Gad2* (**B**). Input (top track) and genomic conservation (dark blue, bottom) are shown for comparison. **C)** Relative fold-change in gene expression for *Pdzrn3* and *Gad2* analyzed by RT-PCR in the MGE of E13 WT (green) and KO (purple) embryos shows upregulation of both genes in KO mice (experiments performed on pools of 3 embryos each, from 3 different litters; n = 6 for WT, n = 4 for KO). **D, F)** Images from *in situ* hybridization experiments for *Pdzrn3* (**D**) or *Gad2* (**F**) transcripts (green) in the MGE of WT and KO embryos at E13, counterstained with DAPI (blue). Dotted lines indicate the border between VZ and SVZ/MZ. Dotted boxes are shown magnified on the right, and show example 100 x 100 *µ*m images from the VZ and SVZ/MZ, as indicated. Scale bars, 100 *µ*m. **E, G)** Quantification of the average number of *Pdzrn3* (**E**) or *Gad2* (**G**) RNA puncta per 100 x 100 *µ*m field of view, as obtained by *in situ* hybridization in the VZ and SVZ/MZ of WT and KO embryos at E13 (n = 4 embryos for both WT and KO). Black bars indicate the mean. Multiple t-tests (panels **C, E**, **G**) were performed; p-values are indicated above the corresponding compared sets of data: those highlighted in bold represent significant differences (p<0.05).

## DISCUSSION

PRDM16 is expressed in radial glia in all telencephalic proliferative zones, suggesting that it might play a key role in the specification of most forebrain neuron lineages (Baizabal et al., 2018; Chuikov et al., 2010; Inoue et al., 2017; Shimada et al., 2017). It is unclear whether *Prdm16* regulates the same progenitor cell behaviors and transcriptional programs in dorsal and ventral progenitors. In the cortex, PRDM16 regulates developmental enhancers involved in the specification and migration of upper layer pyramidal neurons by promoting indirect neurogenic divisions of RG and producing transit-amplifying intermediate progenitors (Baizabal et al., 2018). Here we show that loss of *Prdm16* in MGE progenitors leads to decreased proliferation and reduced interneuron numbers in the cortex and hippocampus; however, in contrast to the cortical *Prdm16* mutant phenotype, this reduction is not layer- or neuronal subtype-specific (**Figures 1** **and S1**). The decrease in MGE-derived interneurons is not due to an increased rate of developmental cell death within this population (**Figure S3**), but rather to defects in the proliferation of *Nkx2.1*-expressing progenitors (**Figure 4**). We observed both a reduction in the number of dividing progenitors and a decrease in their proliferative capacity, as evidenced by the smaller size of retrovirus-labeled clones in the mutant MGE; this might be due to a number of reasons, including alterations in the cell cycle dynamics (such as cell cycle length), changes in the division mode of radial glia, and/or the presence of different types of progenitors in the mutant MGE (Glickstein et al., 2009; Petros et al., 2015; Pilz et al., 2013; Ross, 2011). The loss of both early and late born neuronal lineages from the MGE could reflect key differences in the general role that transit-amplifying progenitors might play in different proliferative regions: whereas in the cortex they have been shown to be the main source of upper-layer pyramidal neurons (Mihalas et al., 2016), their contribution to the neuronal output of the MGE is still largely unexplored. Different subtypes of transit-amplifying MGE progenitors are biased towards the generation of PV+ and SST+ cortical interneurons (Petros et al., 2015), but it is unknown whether transit-amplifying cells are required throughout the entire neurogenic period. Based on our results, we propose that the majority of MGE-derived interneurons are generated through a transit-amplifying progenitor; the decreased ability of radial glia to transition into this type of progenitor in *Prdm16* mutants is reflected by the uniform loss of interneurons across cortical layers and between both histological subgroups. In the future it will be important to determine the fate potential and diversity of transit-amplifying progenitors throughout the neurogenic period for GABAergic cortical interneurons.

The vast majority of cortical interneurons are derived from two distinct ventral telencephalic sources, the MGE and CGE (Wonders and Anderson, 2006). In our study, the loss of MGE-derived interneurons in mutant cortices appears to be partially compensated by an increase in the number of interneurons from a CGE-derived population (**Figure 2**). This is in line with previous research (Denaxa et al., 2018), suggesting a homeostatic mechanism for setting interneuron numbers in response to network activity within the developing cortex. We tested for the first time the capacity of these compensatory cells to restore inhibitory tone. Our electrophysiological recordings demonstrate that a compensatory increase in the number of CGE-derived interneurons was not enough to restore inhibitory inputs onto cortical pyramidal cells to wildtype levels (**Figure 3**). This is likely due to the specific circuit and functional features of Reelin+ interneurons which cannot fill in for the inhibitory circuit functions of PV and SST subgroups derived from the MGE (Olah et al., 2007; Pesold et al., 1999). This suggests that maintaining the proper number and diverse complement of cortical interneuron subgroups derived from different developmental sources is necessary to maintain inhibitory balance (Denaxa et al., 2018).

In both the MGE and the developing cortex PRDM16 binds to cis-regulatory elements in the genome and prevents premature neuronal differentiation by repressing neuronal maturation genes (**Figures 5, 6, S4, S5**) (Baizabal et al., 2018). This is consistent with the observation that PRDM16 binds to a largely overlapping set of distal regulatory elements associated with marks of open chromatin in both MGE and cortical progenitors (**Figure 5**). Just as in the cortex, *Prdm16* repressor function in the MGE might serve to ensure proper timing of lineage-specific gene expression programs to ensure timely transitions from radial glia into transit-amplifying progenitors, which in turn would be responsible for the generation of sufficient numbers of cortical interneurons. While many common aspects of cortical and MGE progenitor function likely require overlapping networks of genes, there are fundamental differences in the migratory capacity and neurotransmitter identity of their neuronal progeny. How might PRDM16 contribute to lineage-specific differentiation programs? In the case of genes that are regulated by PRDM16 in both cortical and MGE progenitors, such as *Pdzrn3*, which encodes an E3 ubiquitin ligase-RING domain containing protein, the molecular function of the encoded protein could be conserved, but the cellular response might be different, eliciting distinct effects on cell fate determination in each proliferative region.

It is also likely that PRDM16 regulates lineage-specific differentiation programs through associations to genomic loci that are exclusive to MGE progenitors. Indeed, one of the MGE-exclusive target genes regulated by PRDM16 is *Gad2*, a gene that is necessary for the synthesis of the inhibitory neurotransmitter GABA (Erlander et al., 1991). It is notable that MGE-specific PRDM16-bound genomic regions are enriched for sequence motifs of basic helix-loop-helix family transcription factors such as ASCL1, which is expressed exclusively in ventral telencephalic neural progenitors and known to regulate GABAergic neuron specification (Long et al., 2009). Many bHLH transcription factors such as OLIG2, ASCL1, and NeuroD family members are expressed exclusively in either ventral or dorsal progenitors (Casarosa et al., 1999; Lu et al., 2000; Osorio et al., 2010). Prdm family members have been shown to interact with basic helix-loop-helix (bHLH) transcription factors (Hohenauer and Moore, 2012; Kinameri et al., 2008; Ross et al., 2012), which are in many cases critical for proper neuronal production by neural progenitors, both in the dorsal and the ventral telencephalon. It has been proposed that interactions between Prdm family proteins and bHLH transcription factors is a conserved mechanism for the regulation of gene expression during neural development (Hohenauer and Moore, 2012; Ross et al., 2012). Such transcription factor complexes may function in the proliferative zones where they are expressed in order to direct differentiation/lineage progression programs towards region-specific neuronal subtype specification (Lindtner et al., 2019). Our genomic analysis of PRDM16 binding in MGE progenitors sheds light on the lineage-specific transcriptional programs regulated by PRDM16 and suggests that specialized PRDM16-associated protein complexes may orchestrate these programs in different progenitor types.

In humans the *PRDM16* gene is located in a distal critical region of chromosome 1 that is deleted in 1p36 deletion syndrome (Jordan et al., 2015). Patients affected by this condition exhibit a spectrum of clinical features that includes intellectual disability, developmental delay and epileptic seizures. It is still unknown to which extent this phenotype is caused by the deletion of PRDM16, but it is likely that distinct mechanisms underlie each of its different clinical aspects. Our study begins to unravel the important question of how PRDM16 controls the fate of progenitors by regulating specific transcriptional programs necessary for neuronal fate determination and excitatory and inhibitory circuit development.

## ACKNOWLEDGMENTS

The authors would like to thank Bernardo Sabatini for providing equipment and resources for the electrophysiology experiments; Mahmoud El-Rifai and Aurélien Begué at the Neurobiology Imaging Facility at Harvard Medical School for the RNAscope experiments; Steve Vu for technical support; all members of the Harwell, Segal and Goodrich labs for discussions and comments; and A. Denise Garcia, Rebecca Ihrie, and Debby Silver for their comments on the manuscript. M.T.G. was partially supported by the Ellen R. and Melvin J. Gordon Center for the Cure and Treatment of Paralysis. Research in C.H.’s laboratory is supported by NIH Grants R56MH119156 and R01NS102228.

## AUTHOR CONTRIBUTIONS

Conceptualization, M.T.G. and C.C.H.; Investigation, M.T.G., J-M.B., D.T., R.P., W.W., Y.X., S.B.; Formal Analysis, M.A.A., M.A.B and M.Y.T.; Resources, C.C.H.; Writing – original draft, M.T.G. and C.C.H.; Writing – review & editing, M.T.G., J-M.B., D.T., R.P., Y.X., M.A.A., C.C.H.; Funding Acquisition, C.C.H.; Supervision, C.C.H.

## RESOURCES TABLE

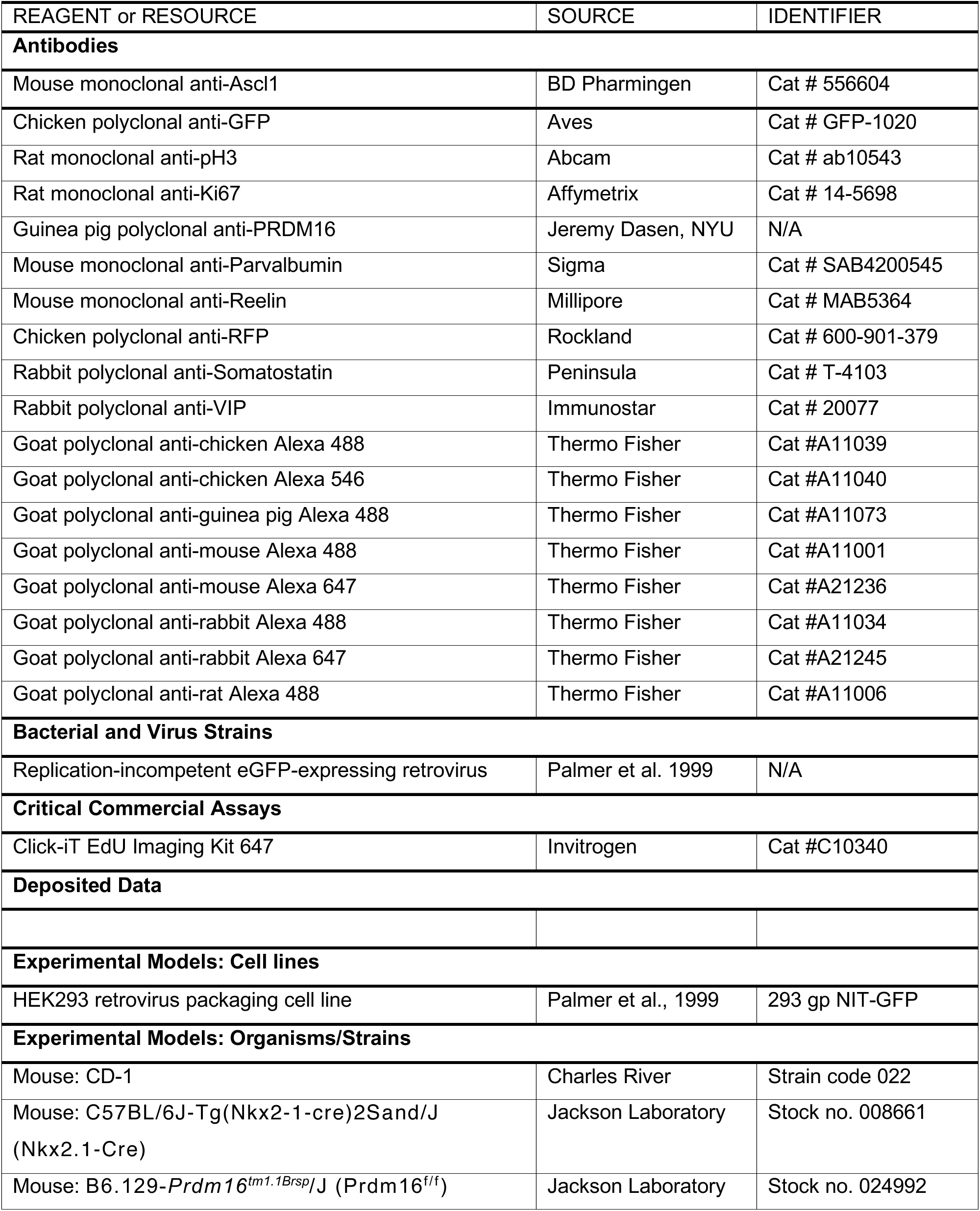
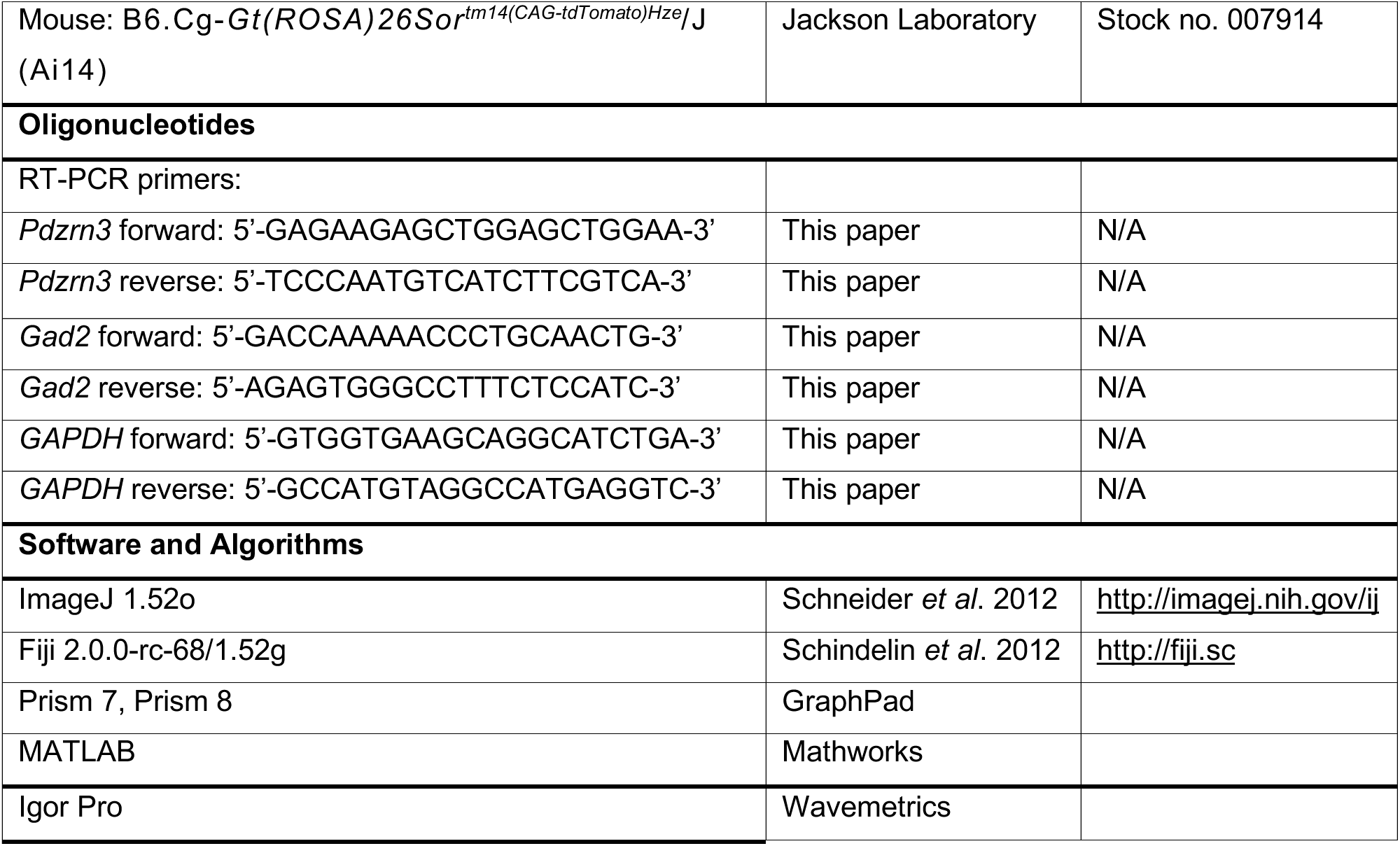

## LEAD CONTACT AND MATERIALS AVAILABILITY

This study did not generate new unique reagents. Further information and requests for resources and reagents should be directed to and will be fulfilled by the Lead Contact, Corey Harwell.

## EXPERIMENTAL MODEL AND SUBJECT DETAILS

All animal procedures conducted in this study followed experimental protocols approved by the Institutional Animal Care and Use Committee of Harvard Medical School. Mouse lines are listed in the Resources Table. Mouse housing and husbandry conditions were performed in accordance with the standards of the Harvard Medical School Center of Comparative Medicine. Mice were group housed in a 12 h light/dark cycle, with access to food and water *ad libitum*. Samples were obtained from animals at embryonic days (E)13, E14, E17, and postnatal days (P)0, P7, P14, P21 and P30, as indicated in Figure Legends. All results reported in adult (P30) and late postnatal stages (P14, P21) include animals of both sexes; the sex of embryos and animals at early postnatal stages was not determined.

## METHOD DETAILS

### Tissue processing

Postnatal animals were transcardially perfused with PBS followed by 4 % paraformaldehyde (PFA) in 120 mM phosphate buffer; their brains were dissected out and post-fixed in 4 % PFA for 2-4 h at room temperature or overnight at 4 °C. Brains were sectioned into 75-100 *µ*m sections on a vibratome (Leica Microsystems VT1200S) and either further processed for immunofluorescence staining or stored at 4 °C in PBS with 0.05% sodium azide. Embryonic brains were dissected out in ice-cold PBS and fixed in 4 % PFA for 2-4 h at room temperature or overnight at 4 °C. Their brains were cryoprotected in 30 % sucrose/PBS overnight at 4 °C, embedded in O.C.T. compound (Sakura), frozen, and stored at −20 °C. Samples were sectioned at 20 *µ*m on a cryostat (Thermo Fisher CryoStar NX70); sections were either stored at −20 °C or further processed for immunofluorescence staining. For fresh frozen samples (used for FISH, see below), the brains were extracted in PBS and immediately embedded in O.C.T. compound, then frozen on dry ice.

### Immunofluorescence staining

Floating vibratome sections: samples were permeabilized with 0.5 % Triton X-100 (Amresco) in PBS for 1-2 h and blocked in blocking buffer (10 % goat serum, 0.1 % Triton X-100 in PBS) for 1-2 h at room temperature. The sections were then incubated for 24-72 h with primary antibodies diluted in blocking buffer at 4 °C. The samples were washed three times (10-30 min/wash) with PBS, counterstained with DAPI (4’,6-diamidino-phenylindole, Invitrogen) for 45 min (both steps at room temperature), and incubated with secondary antibodies diluted in blocking buffer for 2 h at room temperature or overnight at 4 °C. They were then washed (three 10-30 min washes) and mounted on slides with ProLong Gold Antifade Mountant (Invitrogen). Cryosections: slides were allowed to reach room temperature, and then washed three times with PBS. Sections were permeabilized with 0.5 % Triton X-100 in PBS for 30 min, and blocked with blocking buffer for 1 h at room temperature. Slides were incubated with primary antibodies diluted in blocking buffer overnight, in a humid chamber at 4 °C. They were then washed with PBS (three 10-30 min washes), counterstained with DAPI (45 min), and incubated for 1-2 h with secondary antibodies diluted in blocking buffer, at room temperature. Slides were washed (three 10-30 min washes) with PBS and mounted with ProLong Gold Antifade Mountant.

### Imaging and image analysis

Images were acquired using either a Leica DM5500B wide field microscope or a Leica SP8 laser point scanning confocal microscope. 10x, 20x and 25x objectives were used, and the parameters of image acquisition (speed, resolution, averaging, zoom, z-stack, etc.) were adjusted for each set of samples. Images were further analyzed using ImageJ, both in its native and Fiji distributions, as described below. Brightness and contrast were adjusted as necessary for visualization, but the source images were kept unmodified.

### Acute slice preparation and electrophysiology

Mice (19-21 days old) were anesthetized by isoflurane inhalation and perfused transcardially with ice-cold artificial cerebrospinal fluid (ACSF) containing (in mM): 125 NaCl, 2.5 KCl, 25 NaHCO_3_, 2 CaCl_2_, 1 MgCl_2_, 1.25 NaH_2_PO_4_ and 25 glucose, 310 mOsm per kg. Cerebral hemispheres were removed and sliced in cold ACSF (300 *µ*m coronal slices on a Leica VT1200S vibratome). Coronal slices containing somatosensory cortex were recovered for 15-20 min at 34 °C in choline-based recovery solution (in mM): 110 choline chloride, 25 NaHCO_3_, 2.5 KCl, 7 MgCl_2_, 0.5 CaCl_2_, 1.25 NaH_2_PO_4_, 25 glucose, 11.6 ascorbic acid, and 3.1 pyruvic acid, and then transferred to a holding chamber with 34 °C ACSF that progressively cooled down to room temperature (20–22 °C). All recordings were obtained within 1-6 h after slicing and solutions were constantly bubbled with 95% O_2_/5% CO_2_. Individual slices were transferred to a recording chamber mounted on an upright microscope (Olympus BX51WI) and continuously perfused (1–2 ml/min) with ACSF at room temperature. Cells were visualized using a 40× water-immersion objective with infrared DIC optics. Whole-cell voltage-clamp recordings (room temperature) were made from pyramidal cells in L2/3 somatosensory cortex. Patch pipettes (2–4 MΩ) pulled from borosilicate glass (BF150-86-7.5, Sutter Instruments) were filled with a Cs^+^-based internal solution containing (in mM) 130 CsMeSO_4_, 10 HEPES, 1.8 MgCl_2_, 4 Na_2_ATP, 0.3 NaGTP, 8 Na_2_-phosphocreatine, 10 CsCl_2_, and 3.3 QX-314 (Cl^−^ salt), pH 7.3 adjusted with CsOH; 295 mOsm per kg. For all voltage-clamp experiments, errors due to voltage drop across the series resistance (<20 MOhm) were left uncompensated. To isolate mIPSCs, cells were held at −0 mV and ACSF included 20 μM NBQX (Tocris), 10 μM (R)-CPP (Tocris) and 1 μM Tetrodotoxin (Sigma). Membrane currents were amplified and low-pass filtered at 3 kHz using a Multiclamp 700B amplifier (Molecular Devices), digitized at 10 kHz and acquired using National Instruments acquisition boards and a custom version of ScanImage written in MATLAB (Mathworks). Off-line analysis of mIPSC frequency was performed using custom routines written in MATLAB and Igor Pro (Wavemetrics). Statistical analyses were done in GraphPad Prism 7 software (GraphPad).

### Viral production and *in utero* injection

The retrovirus packaging cell line HEK293 gp NIT-GFP was grown to 90 % confluence in DMEM supplemented with 10 % fetal bovine serum. Cells were transfected with a pCMV-VSV-G vector using Lipofectamin 2000 (Invitrogen) and Optimem (GIBCO). Two days after transfection, the cell supernatant was collected, filtered through a 0.45-*µ*m filter (VWR International) and centrifuged at 25000 rpm for 90 min at 4 °C. After discarding the supernatant, 100 *µ*l of cold PBS+Ca^2+^ were added to the pellet, which was incubated overnight at 4 °C. Viral particles were then resuspended, aliquoted, and stored at −80 °C. The titer of the viral preparation was determined to be 10^6^-10^7^ cfu. Timed pregnant mice were anesthetized using an isoflurane vaporizer and placed on a warming pad. An abdominal incision (approx. 2 cm wide) was made, and the uterine horns were exposed on top of a sterile gauze pad. The uterus was periodically moistened with sterile PBS pre-warmed at 37 °C during the entire procedure. Approximately 0.5-1 *µ*l of retrovirus, mixed with 0.05 % Fast Green for visualization was injected into the lateral ventricles of each embryo, using heat-pulled beveled glass micropipettes (Drummond). The abdominal cavity was sutured and stapled before administering buprenorphine (0.05-0.1 mg/kg) and ketoprofen (5-10 mg/kg). The pregnant dams were allowed to recover in a 35 °C chamber for 1-2 h after surgery, then returned to usual housing conditions and sacrificed 24 h after surgery; the embryos were retrieved and processed as explained above.

### Chromatin immunoprecipitation and sequencing

The MGE of 18 embryos from 2 litters of E14 wildtype mice were dissected out and finely minced. Cells were dual crosslinked by incubating in 1.5 mM EGS (ethylene glycol bis[succinimidyl succinate]) solution for 20 min with rotation, and then with 1 % PFA + 1.5 mM EGS for an additional 10 min (both steps were performed at room temperature). Crosslinking was quenched by adding glycine to a final concentration of 125 mM and rotating for 5 min at room temperature. Cells were then washed twice with cold PBS + EDTA-free protease inhibitor, centrifuged and stored at −80 °C or freshly resuspended in lysis buffer (20 mM Tris-HCl pH 8.0, 85 mM KCl, 0.5 % NP40) and incubated on ice for 30 min. Nuclei were pelleted by centrifugation at 1500 *g* for 5 min, resuspended in SDS buffer (0.2 % SDS, 20 mM Tris-HCl pH 8.0, 1 mM EDTA) and incubated on ice for 10 min. Nuclei were then sonicated using a Covaris S2 ultrasonicator for shearing chromatin into fragments with a size range of 100-500 bp. After spinning chromatin at 18,000 *g* for 10 min, supernatant was transferred to a clean tube and one volume of 2X ChIP dilution buffer (0.1 % sodium deoxycholate, 2 % Triton X-100, 2 mM EDTA, 30 mM Tris-HCl pH 8.0, 300 mM NaCl) was added. At this step, a volume of supernatant containing around 0.5 million nuclei was set aside as input control and the remaining supernatant was incubated with 5 mg of anti-PRDM16 antibody overnight at 4 °C with rotation. Next day, 50 ml of washed protein G beads (22.5 mg/ml; Novex) were added to the chromatin solution and incubated for 2 h at 4 °C. After incubation, beads were washed twice with low salt wash buffer (0.1% SDS, 1% Triton X-100, 2 mM EDTA, 20 mM Tris-HCl pH 8.0, 150 mM NaCl) followed by two washes with high salt wash buffer (0. 1 % SDS, 1 % Triton X-100, 2 mM EDTA, 20 mM Tris-HCl pH 8.0, 500 mM NaCl) then two washes with LiCl wash buffer (0.25 M LiCl, 0.5 % NP40, 0.5 % sodium deoxycholate, 1 mM EDTA, 10 mM Tris-HCl pH 8.0) and finally two washes with TE pH 8.0 (10 mM Tris-HCl, 1 mM EDTA). Beads were then resuspended in 90 ml of freshly prepared ChIP elution buffer (1 % SDS, 0.1 M NaHCO3) and incubated at 65 °C for 30 min with rotation. The recovered supernatant was incubated in reverse crosslinking solution (250 mM Tris-HCl pH 6.5, 62.5 mM EDTA pH 8.0, 1.25 M NaCl, 5 mg/ml of Proteinase K) at 65 °C overnight. DNA was then extracted with phenol/chloroform/isoamyl alcohol, precipitated with 3 M sodium acetate pH 5.0 and resuspended in TE pH 8.0 low EDTA (10 mM Tris-HCl, 0.1 mM EDTA). Finally, samples were treated with RNase A (100 mg/ml) for 30 min at 37 °C. For library preparation, genomic DNA was purified, end repaired, ligated with barcoded adaptors, amplified for 14 PCR cycles and purified using the Ovation Ultralow System V2 (NuGEN) according to manufacturer’s instructions. Library fragments in the range of 100-800 bp were size-selected using agarose gel electrophoresis followed by DNA gel extraction (QIAGEN). Recovered DNA was further cleaned and concentrated using a column (Zymo Research). Libraries were sequenced in an Illumina HiSeq 2500 sequencer to a sequencing depth of 30-40 million reads per sample.

### ChIP-Seq analysis

Reference genomic sequence and annotations for the mouse genome (mm10) were obtained from the University of California Santa Cruz Genomics Institute (Casper et al, 2018). ChIP-Seq reads generated from PRDM16 ChIP and input libraries were aligned to the mouse genome using bowtie2 (Langmead et al., 2012) with default parameters except for “-I 50 -X 750”. The read densities were calculated with the Gaussian kernel (bandwidth of 35 bp) using the R package SPP (Kharchenko et al., 2008). Peaks were called using MACS2 (version 2.1.2) (Zhang et al., 2008) with “callpeak” and a p-value threshold of 0.05 followed by Irreproducible Discovery Rate (IDR) analysis (as described by Landt et al., 2012) with a threshold of <= 0.05. The MGE peak set was compared to the previously published cortical PRDM16 ChIP experiment (Baizabal et al.) with a merged set of peaks defined by the union of the MGE and cortical peak sets. Peaks were considered to be present in both cortical and MGE samples if the ChIP/input ratio of library-size normalized read counts within the peak region was >= 3 for both experiments. Peaks were considered “cortical only” (“MGE only”) if ChIP/input ratio was >=3 for the cortical (MGE) experiment and <= 1.5 for the MGE (cortical) experiment. Gene ontology enrichment analysis was performed using the R package ClusterProfiler (Yu et al., 2012) with the parameters ont=“BP” and maxGSSize=20000.

Motif Analysis was performed on set peaks detected in the MGE using HOMER (Heinz et al., 2010). *De novo* motif analysis was conducted by parsing input sequences at +/- 100 bp from peak center. The FDR-corrected probability of that motif being over-represented amongst target sequences is presented as a q-value known motif enrichment is screened against the JASPAR database of previously determined high quality motifs.

### Quantitative reverse transcriptase polymerase chain reaction (qRT-PCR)

The MGE of E13 WT and KO embryos were dissected out in PBS and stored at −80 °C. Total RNA was extracted using the Qiagen RNeasy Mini Kit. First strand synthesis was performed using Invitrogen SuperScript III First-Strand Synthesis kit with Oligo(dT)_20_ primers. Quantitative PCR was performed using Sybr Green Supermix (BIO-RAD) on a C1000 Touch thermocycler (BIO-RAD). Primers designed for each tested gene can be found in the Key Resources Table. Relative levels of mRNA were calculated using the delta CT method with *Gapdh* as the internal control.

### Fluorescent in situ hybridization (FISH)

Fresh frozen samples (see above) were submitted to the RNAscope protocol (Advanced Cell Diagnostics), following the manufacturers’ instructions. RNAscope probes (see Resources Table) against *Pdzrn3* and *Gad2* were used in combination with a *tdTomato* probe to confirm that all areas analyzed were within the *Nkx2.1*-expressing portion of the MGE.

## QUANTIFICATION AND STATISTICAL ANALYSIS

### Cell quantification

The CellCounter tool in ImageJ/Fiji was used for all cell quantifications. In the mature cortex (Figures 1C-J, 2A-J), 1 mm-wide sections were selected from images obtained at the level of the somatosensory cortex. All cells positive for the markers analyzed in each case were quantified within each cortical layer, as identified by the distribution of DAPI-counterstained nuclei. In the hippocampus (Figure S1), the entire structure was analyzed at equivalent rostro-caudal levels; layers and areas were determined by the distribution of DAPI-counterstained nuclei. In the developing cortex at different stages (Figure 3A), the entire surface of the neocortex was measured on both hemispheres of each sample at equivalent rostro-caudal levels, and the number of cells positive for CC3 and tdT was quantified within the measured surface, then normalized by mm^2^. In E13 MGE samples (Figure 3B), the entire surface of the MGE in sections at equivalent rostro-caudal locations was measured; cells positive for pH3 were quantified within the measured area and normalized by mm^2^. For the quantification of MGE clones (Figures 3D-F), cells were considered as part of the same clone if they had similar fluorescence intensities and were located within 50 *µ*m of the radial fiber in the case of RG-containing clones or within a 50-*µ*m radius in the case of IP-containing clones. Clones were not analyzed if other GFP-labeled cells of different fluorescence intensity were present in their vicinity (within 100 *µ*m of the radial fiber / within a 100-*µ*m radius). Quantification of RNA puncta per unit area was performed using an automated data processing pipeline in MatLab, guided by MatBots (https://hms-idac.github.io/MatBots); each data point (Figure 6E, G) corresponds to the average values from the analysis of three 100 x 100 *µ*m fields selected within the VZ and SVZ/MZ of two sections/MGEs per sample (except for one KO sample, where only one section was analyzed). Cell and RNA puncta numbers were compiled in Microsoft Excel spreadsheets; GraphPad Prism 8 (GraphPad) was used to build graphs.

### Statistical analysis

All statistical analyses were performed with GraphPad Prism 8, as detailed in the Figure Legends. All p-values were rounded to ten thousandth, and are presented above each statistical comparison in the corresponding figures; those highlighted in bold are below 0.05, which was considered the cutoff for statistical significance (p-values deemed not statistically significant under this criterion are displayed in parenthesis above the corresponding comparisons in the Figures).

## SUPPLEMENTARY FIGURE LEGENDS

**Figure S1, related to Figure 1. Deletion of *Prdm16* in the *Nkx2.1* lineage causes loss of hippocampal interneurons. A)** Representative images of the hippocampus of WT and KO mice at P30, after immunofluorescence staining for tdTomato (red), counterstained with DAPI (blue). Scale bars, 500 *µ*m. Area labels (WT image): CA – *cornus Ammonis* (Ammon’s horn) regions 1, 2 and 3; DG – dentate gyrus; S – subiculum. Layer labels (KO image): so – *stratum oriens*; sp – *stratum pyramidale*; sr – *stratum radiatum*; slm – *stratum lacunosum moleculare*; sl – *stratum lucidum*; ml – molecular layer; g – granule cell layer; h – hilus. **B)** Quantification of the number of tdTomato+ cells in the entire hippocampus of WT (green squares) and KO (purple circles) mice (n = 10 for WT, n = 12 for KO). **C)** Quantification of the total number of tdTomato+ cells across different areas of the entire hippocampus in WT and KO mice at P30. **D)** Quantification of the number of tdTomato+ cells across different cellular layers in the entire hippocampus of WT and KO mice at P30. **E)** Quantification of the total number of somatostatin+ cells in the entire hippocampus of P30 WT (n = 4) and KO (n = 5) mice. **F)** Quantification of the total number of somatostatin+ cells in the entire hippocampus of P30 WT (n = 4) and KO (n = 5) mice. **G, H)** Analysis of the distribution of tdTomato+ cells costained for SST (**G**) or PV (**H**) across the indicated layers of the hippocampus of P30 WT and KO mice in areas CA1, CA2, CA3 and DG. Means ± S.D. are represented. Unpaired t-tests with Welch’s correction (panels **B**, **E**, **F**) or multiple t-tests (panels **C**, **D**) were performed; p-values are indicated above the corresponding compared sets of data: those highlighted in bold represent significant differences (p<0.05).

**Figure S2, related to Figure 2. Partial compensation of depleted MGE-derived interneurons is restricted to the reelin+ population in upper layers. A)** Number of VIP+ cells in each indicated cortical layer, per 1 mm-wide column. **B)** Number of reelin+ cells in each indicated cortical layer, per 1 mm-wide column. **C)** Percentage of VIP+ cells costained for tdTomato. Multiple t-tests (panels **A** and **B**) or unpaired t-tests with Welch’s correction (panel **C**) were performed; p-values are indicated above the corresponding compared sets of data: those highlighted in bold represent significant differences (p<0.05).

**Figure S3, related to Figure 4. Loss of cortical interneurons is not due to increased cell death or defects in migration. A)** Quantification of the number of cells costained with tdTomato (tdT+) and the apoptotic marker cleaved caspase 3 (CC3+) per mm^2^ in the cortex of WT (green circles) and KO (purple squares) mice at the indicated developmental stages. Error bars represent S.D. Multiple t-tests were performed; p-values are indicated above the corresponding compared sets of data: those highlighted in bold represent significant differences (p<0.05). **B)** Overview of the migration of Nkx2.1-lineage cells (expressing tdTomato, white) within coronal hemisections of WT and KO brains at E11, E13 and E15. Empty arrowheads indicate the extent of migration of MGE-derived interneurons into the cortex within each section. Scale bars, 500 *µ*m.

**Figure S4, related to Figure 5. Comparison of PRDM16 ChIP-Seq peaks between cortex and MGE. A)** Distribution of E15 cortex PRDM16 ChIP-Seq peaks across the genome (TSS: transcription start site; TTS: transcription termination site). **B)** Top 30 gene ontology term enrichment in genes closest to PRDM16 ChIP-Seq peaks from E14 MGE (this study) and E15 cortex (Baizabal *et al*. 2017) experiments. Categories represent all PRDM16 peaks from both sets of experiments (‘All’), common to both datasets (‘Both’), or exclusive to either the cortex (‘Cortical Only’) or the MGE (‘MGE Only’).

**Figure S5, related to Figure 5. Analysis of PRDM16 peaks relative to their genomic location. A)** Distribution of E13 MGE PRDM16 ChIP-Seq peaks across the genome (TSS: transcription start site; TTS: transcription termination site) – note: this panel is the same data displayed as Figure 5C, and is included here just for clarity. **B)** Top 30 gene ontology term enrichment in genes closest to PRDM16 ChIP-Seq peaks located in TSS, intronic or intergenic locations, as indicated. **C-E)** Read density (fragments per million reads) in several embryonic ChIP-Seq experiments, within a genomic window centered around PRDM16 binding sites (left), and read density alignment of the ChIP-Seq experiments (right), analyzed by the location of the peaks in either TSS (**C**), intronic (**D**) or intergenic (**E**) genomic regions. Data were generated for this study from E14 MGE (dark blue), or obtained from Baizabal *et al*. 2017 (CTX, ChIP-Seq for PRDM16 in E15 cortex; light blue) and Sandberg *et al*. 2016 (ChIP-Seq for each specified histone modification in E13 MGE; all other plots)

